# Apolipoprotein E interacts with amyloid-β oligomers via positively cooperative multivalent binding

**DOI:** 10.1101/473892

**Authors:** S. Ghosh, T. B. Sil, S. Dolai, K. Garai

## Abstract

Interaction of apolipoprotein E (apoE) isoforms with amyloid-β (Aβ) peptides is considered a critical determinant of the progression of Alzheimer’s disease. However, molecular mechanism of the apoE-Aβ interaction is poorly understood. Here we characterize the nature of the apoE-Aβ complexes and identify the region of apoE that interacts with Aβ. We have prepared three distinct fragments of apoE4, *viz.*, the N-terminal fragment (NTF), hinge domain fragment (HDF) and C-terminal fragment (CTF) to compare its interactions with Aβ. Kinetics of aggregation of Aβ is delayed dramatically in presence of low, substoichiometric concentrations of both NTF and CTF in lipid-free, as well as, in lipidated forms. Effect of HDF is found to be small. Strong inhibition by NTF and CTF at substoichiometric concentrations indicate interactions with the ‘intermediates’ or the oligomers of Aβ. Kinetics of Forster Resonance Energy Transfer (FRET) between full-length apoE4 labeled with EDANS at positions 62, 139, 210, 247, and 276 and tetramethylrhodamine (TMR)-labeled Aβ further support involvement of multiple regions of apoE in the interactions. Since the interactions involve intermediates of Aβ quantitative evaluation of the binding affinities are not feasible. Hence we employed a competitive binding assay to examine whether the N- and C-terminal domains interact cooperatively. Addition of unlabeled full-length apoE eliminates the FRET between EDANS-NTF + EDANS-CTF and TMR-Aβ almost completely but not vice versa. Furthermore, full-length apoE but not the equimolar mixture of the fragments could displace the already bound EDANS-apoE molecules from the complexes. Therefore, binding affinity of the Aβ oligomers to the intact full-length apoE is much higher than the affinity to the domains when mixed together as fragments. Thus, our results indicate that apoE-Aβ complex formation is mediated by positively cooperative multivalent binding between the multiple sites on apoE and the oligomeric forms of Aβ.

## Introduction

Pathology of Alzheimer’s disease (AD) is characterized by deposition of amyloid β (Aβ) peptides in the form of senile plaques in the brain. However, the strongest genetic risk factor for AD is the ʵ4 isoform of Apolipoprotein E (apoE) (1). ApoE is a 299 -residue lipoprotein with three major isoforms, *viz.*, apoE2 (Cys 112, Cys 158), apoE3 (Cys 112, Arg 158) and apoE4 (Arg112, Arg 158). The three isoforms differ by single amino acid substitutions, but their effects on the outcome of AD are immense. How the apoE isoforms influence progression and/or onset of the pathology of AD is still not fully clear. However, numerous studies, both *in vitro* and *in vivo*, suggest isoform specific role of apoE proteins on the metabolism of Aβ. For example, apoE4 is found to be associated with impaired uptake, clearance and degradation of Aβ (2-5). Furthermore, presence of the ε4 allele is associated with the higher load of amyloid plaques in the brains of both humans and transgenic mice expressing familial AD mutations (6-8). Thus, interactions of the apoE isoforms with Aβ is considered to be one of the most critical determinants of development of AD.

*In vitro*, apoE isoforms are found to interact with Aβ and alter the kinetics of amyloid aggregation. However, in this regard different groups have reported considerably diverse and sometimes even contradictory observations. For example, a few early studies suggested that apoE isoforms accelerate the aggregation of Aβ in an isoform dependent manner, with apoE4 showing the strongest effects. These studies indicated apoE4 as a pathological chaperone (9, 10). However, most of the recent studies suggest that apoE isoforms delay the kinetics of the fibrillization of Aβ, possibly by stabilizing the soluble oligomers of Aβ. ApoE4 has the highest and apoE2 has the lowest effects (11-13). These authors hypothesized that apoE4 may influence the pathology of AD by increasing the concentration of the soluble oligomers, which are believed to be the major cytotoxic species in AD (11, 14, 15). The apparently contradictory results are believed to arise due to the transient and heterogeneous nature of both apoE and Aβ. For example, oligomers of Aβ are known to be highly heterogeneous and transient (16). *In vivo*, apoE isoforms exist primarily in the lipidated forms, although it is found in lipid-poor and in lipid-free forms as well (17, 18). Furthermore, lipid-free apoE exists as a mixture of monomers, dimers, tetramers and higher order oligomers (19-22). While most experiments confirm direct interactions between apoE and Aβ (11, 23-26), very little is understood about the biophysical properties of these complexes (reviewed in Tai et al) (27).

In this article, we attempt to identify the region of apoE involved in the interactions with Aβ. The structure of apoE consists of three distinct domains, *viz.*, an N terminal domain (residues1-167) which contains the receptor binding regions, a C-terminal domain (residues 238-299), which is responsible for oligomerization and binding to lipids, and a flexible hinge region (residues 168-237), which connects the N- and the C-terminal domains (28). The function of the hinge domain is unknown (22). The N-terminal domain consists of a four helix bundle, while the C-terminal domain contains a major helical region (residues 238-264) and several stretches of short helices and unstructured regions (28). There are considerable confusions in the literature regarding which regions of apoE bind to Aβ. Studies examining apoE-Aβ complexes by domain specific antibodies (25, 29) and intermolecular Forster resonance energy transfer (FRET) (30) suggest that the C-terminal domain of apoE is the primary region interacting with Aβ. Alternatively, a few studies have shown that the N-terminal domain harbors the primary Aβ binding sites (13, 31). A recent study using mass spectroscopy following crosslinking of the interacting species concluded that both the domains of apoE interact with Aβ (32). The apparent confusions in the results from these studies arise most likely due to the dynamic and heterogeneous nature of the apoE-Aβ complexes and the possible non-specific interactions of the amyloids with various proteins. The second point relates to the observations that amyloids interact with numerous physiological proteins and sequester these to inclusion bodies possibly via non-specific interactions (33). Therefore, in-depth investigation of the interactions between apoE and Aβ is required to identify the specific interactions from the weak binding interactions. Here we have performed kinetic measurements over long times to delineate the interactions of apoE with the monomers, oligomers and the fibrils of Aβ. We have prepared three different fragments of apoE4, *viz.*, an N-terminal fragment (NTF), a C-terminal fragment (CTF) and a Hinge domain fragment (HDF) to examine the effects of the fragments on the time course of the aggregation of Aβ42. Furthermore, we have prepared fluorescently labeled NTF, CTF and several single cysteine mutants of full-length apoE4 to monitor FRET between the various regions of apoE and Aβ. Finally, we have used a competitive binding assay to examine the relative affinities of the sum of the fragments compared to the intact full-length apoE to Aβ. Our results suggest that all the three fragments of apoE in isolation interact with Aβ. However, the intact form of apoE interacts with the highest affinity.

### Materials and Methods

All the chemicals unless otherwise stated are purchased from Sigma Aldrich (Sigma, St. Louis, MO).

### Purification of Aβ42

Chemically synthesized Aβ42, TMR-labeled Aβ42 (TMR-Aβ42) and EDANS-labeled Aβ42 (EDANS–Aβ42) are purchased from AAPPTec LLC (Louisville, KY, USA). TMR is attached to the N-terminal amine as described previously (34). EDANS is attached to the N-terminal of Aβ using fmoc-Glu(EDANS)-OH. Aggregation properties of TMR-Aβ has been reported earlier (34, 35). Stock solutions of Aβ42, TMR-Aβ42, and EDANS-Aβ42 have been prepared by using the protocol described by Sil et al (34). Briefly, lyophilized powder of the peptide is dissolved in formic acid, flash frozen in liquid nitrogen and lyophilized again. The lyophilized powder is dissolved in 6 M GdnCl containing 10 mM PBS, 1 mM EDTA and 5 mM βMe. The solution is purified further by size exclusion chromatography using a Superdex 75 column (GE Healthcare, USA) in 4 M GdnCl containing 10 mM PBS, 1 mM EDTA and 5 mM βMe. The final stock solution of Aβ is prepared in 5 mM NaOH solution containing 1 mM EDTA and 5 mM β-Mercaptoethanol (βMe) following buffer exchange using a PD10 desalting column (GE Healthcare, USA). The stock solution is then aliquoted into 100 µl vials, flash-frozen in liquid nitrogen and stored at −80 0εC.

### Expression and Purification of the full-length and the fragments of apoE4

The WT-apoE4 plasmid is a kind gift from Dr. Carl Frieden (Washington University, St. Louis). The plasmids for all the fragments (NTF, HDF, and CTF), the single cysteine mutants of full-length apoE4 and apoE fragments used here are prepared using the plasmid of WT-apoE4 by GenScript, USA. The proteins are expressed and purified using the protocol described by Garai et al (12). The final step of the purification is performed using size exclusion chromatography using a Superdex 200 column (GE Healthcare) in phosphate buffered saline (PBS) (pH 7.4) containing 1mM EDTA and 5mM βMe. The HDF of apoE is purified using a Superdex 75 column. Lipidated full-length apoE and the apoE fragments are prepared using the protocol described earlier by Garai et al.(12) Briefly, small unilamellar vesicles (SUV) of 1,2-Dimyristoyl-sn-glycero-3-phosphorylcholine (DMPC) are prepared by extrusion through a 50 nm polycarbonate filter (Avanti Polar lipids, USA). ApoE (~20 µM) is incubated with the SUVs (4 mg/ml) at 25 0εC for overnight (O/N). The mixture is then purified using size exclusion chromatography to remove free lipids as well as unbound apoE using a Superdex 200 column in PBS, pH=7.4, buffer in presence of 1 mM EDTA, and 5 mM βMe. The lipidation of apoE is confirmed using fluorescence spectrum of the tryptophan residues in apoE with a characteristic blue shift of the peak (see Figure S1). We have noticed that lipid-free apoE is prone to oxidation and oligomerization (data not shown). Hence we add 5 mM βMe to all the solutions of apoE.

### Fluorescence labeling of the single cysteine mutants of apoE4 and the fragments

Fluorescence labeling of the single cysteine mutants of full-length and the fragments of apoE4 is performed using standard maleimide labeling protocol (19). Briefly, approximately 2 mg lyophilized powder of the protein is dissolved in 1 ml PBS, pH 7.4 buffer containing 6 M urea and 2 mM Tris(2-carboxyethyl)phosphine (TCEP). The solution is degassed for 30 minutes at room temperature. EDANS-maleimide dye (Anaspec) is then added in 5-fold molar excess to the protein solution and degassed for 3 hours at room temperature. The EDANS-apoE is then purified by size exclusion chromatography using a Superdex 200 column to remove the unbound dye. The labeling efficiency, measured using the ratio of the absorbances at 340 nm and 280 nm, is estimated to be within 80-90% for the different cysteine mutants of apoE.

### Measurement of the time course of amyloid aggregation by ThT or TMR fluorescence

Aggregation of Aβ42 is monitored by Thioflavin T (ThT) fluorescence and the aggregation of TMR-Aβ42 is monitored by TMR fluorescence. The stock solution of Aβ42 (130 µM) is diluted to the final concentration (5 µM) in PBS buffer (pH = 7.4) containing 1 mM EDTA, 5 mM βMe, and 8 µM Thioflavin T (ThT). The full-length or the fragments apoE4 in the lipid-free or in the lipidated states are added to the solution of Aβ42 at t = 0. The samples are incubated in clean glass tubes (i.d. = 10 mm) at 25 0εC in the cuvette holder inside the spectrofluorometer (PTI, USA). The samples are stirred continuously using a 3 mm × 6 mm Teflon coated magnetic bead. The time course of aggregation of Aβ42 is monitored continuously using fluorescence of ThT (λ_ex_ = 438 nm, λ_em_ = 485 nm). The aggregation kinetics of 1 µM TMR-Aβ42 are monitored continuously using fluorescence of TMR (λ_ex_ = 520 nm and λ_em_ = 600 nm) (35). Full-length or the fragments of apoE4 in the lipidated or lipid-free forms are added to this solution at t = 0.

### ApoE-Aβ interactions using Forster Resonance Energy Transfer (FRET)

Direct interactions between apoE and Aβ are monitored using kinetics of FRET between unlabeled apoE and EDANS-Aβ42. The concentrations of apoE4 (or the fragments) and EDANS-Aβ42 used are 500 nM and 5 µM respectively. In these experiments, the tryptophan residues (W20, W26, W34 and W39, W210, W264, and W276) of apoE are used as the FRET donors and the EDANS attached to Aβ as the acceptor. Therefore, the number of tryptophan residues in the NTF, HDF, and CTF are four, one and two respectively. The samples are incubated at 25 0εC with continuous stirring in quartz cuvettes (FireflySci, USA) placed in the temperature-controlled cuvette holder inside the spectrofluorometer. The FRET is monitored using the fluorescence of tryptophan (donor) in apoE with λ_ex_ = 290 nm and λ_em_ = 350 nm. Fluorescence of apoE in presence of unlabeled Aβ42 under the same experimental conditions are used as controls.

To identify the regions of apoE involved in the interactions, FRET experiments are performed using an EDANS-labeled single cysteine mutant of full-length apoE4 and TMR-Aβ42. For these experiments, five single cysteine mutants (A62C, S139C, W210C, A247C, W276C) of apoE4 have been used. EDANS labeled apoE4 (250 nM) is incubated in presence of 1 µM TMR-Aβ or 1 µM unlabeled Aβ42 in clean glass test tubes at 25 εC with continuous stirring inside the spectrofluorometer. Fluorescence of the FRET donor (EDANS) is monitored continuously using λ_ex_ = 350 nm and λ_em_ = 482 nm. Interactions with the fragments are monitored using FRET between EDANS-NTF(A102C) or EDANS-CTF(A247C) and TMR-Aβ42.

### Competitive binding assay to compare affinities of the full-length and fragments of apoE for Aβ

A 250 nM solution of EDANS-apoE4(A247C) or EDANS-NTF(A102C) or EDANS-CTF(A247C) or EDANS-NTF+EDANS-CTF is incubated with 1µM TMR-Aβ42 in clean glass tubes at 25 εC with continuous stirring. For the competition assay 1 µM unlabeled full-length or the fragments of apoE4 are added in this solution at t = 0. FRET is monitored continuously using EDANS (donor) fluorescence. To examine the displacement of labeled apoE4 (or the labeled fragments of apoE4) from the EDANS-apoE/TMR-Aβ complexes, 1µM unlabeled apoE4 (or the fragments) is added after t > 10 hours. FRET is monitored continuously using the fluorescence of EDANS (donor).

## Results

### N-, C-terminal and hinge domain fragments of apoE4 delay the kinetics of aggregation of Aβ42

First, we examine how the different fragments (NTF, HDF, and CTF) of lipid-free apoE affect the kinetics of aggregation of Aβ. In all the experiments reported in this article, we have used the full-length or the fragments of the E4 isoform of apoE and the 1-42 alloform of Aβ. Figure 1, A-E show the effects of apoE4 and its fragments on the kinetics of fibrillization of Aβ monitored by the fluorescence of ThT. Expectedly, the kinetics of aggregation of Aβ is characterized by a clear ‘lag’, growth, and saturation phase. Imaging of the Aβ42 aggregates by AFM confirms the fibrillar morphology (Figure S2A). Figure 1A shows that sub-stoichiometric concentrations of NTF of apoE4 delay the kinetics of the aggregation of Aβ in a concentration dependent manner. At lower concentrations of NTF, the ‘lag’ phase is prolonged but the total extent of fibrillization measured by ThT fluorescence remains almost the same. However, at higher concentrations of

**Figure 1:**
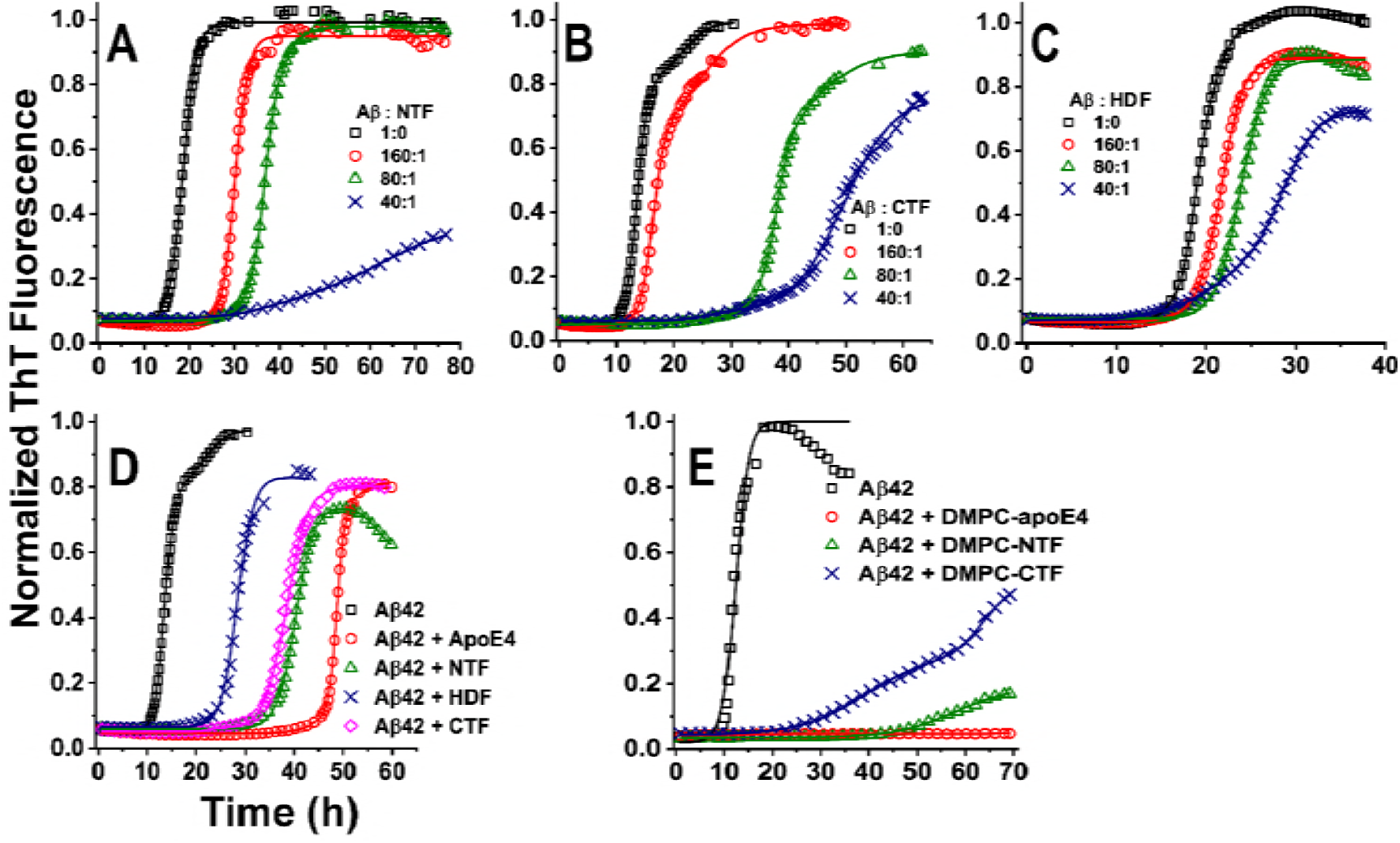
**Fragments and full-length apoE4 delay kinetics of fibrillization of Aβ42.** Panel A-C: aggregation of 5 µM Aβ42 in presence of 0 (□), 31 (○), 62 (Δ) and 125 nM (×) of NTF (A), CTF (B) and HDF (C). Panel D shows comparison of the effects of lipid-free HDF (×), CTF (◊), NTF (Δ) and full-length apoE4 (○). Panel E: Effects of DMPC-CTF (×), DMPC-NTF (Δ) and DMPC-full-length-apoE4 (○). In panels D and E, concentration of Aβ42 and apoE used are 5 µM and 62 nM respectively. The solid lines are guide to the eye. The experiments have been performed in PBS, pH 7.4 buffer at 25 εC. The aggregation is monitored continuously using fluorescence of thioflavin-T (ThT).

NTF, the total fluorescence of ThT is also reduced significantly. Therefore, NTF delays and reduces the fibrillization of Aβ42 in a dose dependent manner. Figure 1B shows that similar effects are also observed in presence of the CTF. Figure 1C shows that HDF also has delaying effects on the aggregation kinetics of Aβ but the effects are significantly less than the NTF and the CTF. Figure 1D compares the effects of the fragments and the full-length apoE4 when used at identical molar concentrations. Clearly, the effect is the highest with the full-length apoE4 followed by the NTF, CTF, and HDF. AFM images of the aggregates indicate that morphology of the Aβ fibrils doesn’t alter significantly in presence of apoE or its fragments (see Figure S2). Taken together the data presented above indicate that both N- and C-terminal fragments of apoE strongly alter the kinetics of aggregation of Aβ. In order to verify if the effects of apoE are specific we have examined the effects of two other unrelated proteins, viz, bovine serum albumin (BSA) and intestinal fatty acid binding protein (IFABP) on the aggregation of Aβ. Supplementary Figure S3 shows that the effects of BSA and IFABP are minimal compared to that of apoE at same molar concentration. Strong effects at substoichiometric concentrations of apoE are consistent with the previously proposed hypothesis that apoE must interact with the ‘intermediates’ or the oligomers of Aβ (11-13, 36). Additionally, considerable effects at low nanomolar concentrations of apoE indicate high affinity of the interactions.

Then we examined how lipidated form of the fragments and the full-length apoE influence aggregation of Aβ. Here lipidation of apoE has been performed using DMPC. Figure 1E shows that DMPC-apoE4, DMPC-NTF and DMPC-CTF all delay the kinetics of aggregation of Aβ. Once again, the effect is the highest for the full-length DMPC-apoE4, followed by the DMPC-NTF and the DMPC-CTF. Taken together, the above results clearly demonstrate that the fragments of apoE both in the lipid-free and in the lipidated forms interact with Aβ to delay the kinetics of fibrillization. It may be noted that lipid-free apoE undergoes large conformational changes upon binding to lipids (37). Why both the forms interact with Aβ with high affinities is not clear. We speculate that this could happen due to dynamic equilibrium between the lipidated and the lipid-free forms of apoE (38). We note here that we haven’t used DMPC-HDF because this HDF couldn’t be lipidated (data not shown).

### Comparison between the full-length intact apoE4 and the fragments mixed together

The data presented above pose an interesting question about whether the combined effects of fragments are comparable to the full-length apoE4. To discern that, we mixed the fragments of apoE4 together and compared its effects with that of the same molar concentration of the full-length apoE. Figure 2A shows that the effects of NTF, CTF, and HDF mixed together are less than that of the full-length apoE4 but the effects are comparable. Similar observations are made in case of DMPC-NTF, DMPC-CTF and the full-length lipidated apoE4 (Figure 2B).

**Figure 2:**
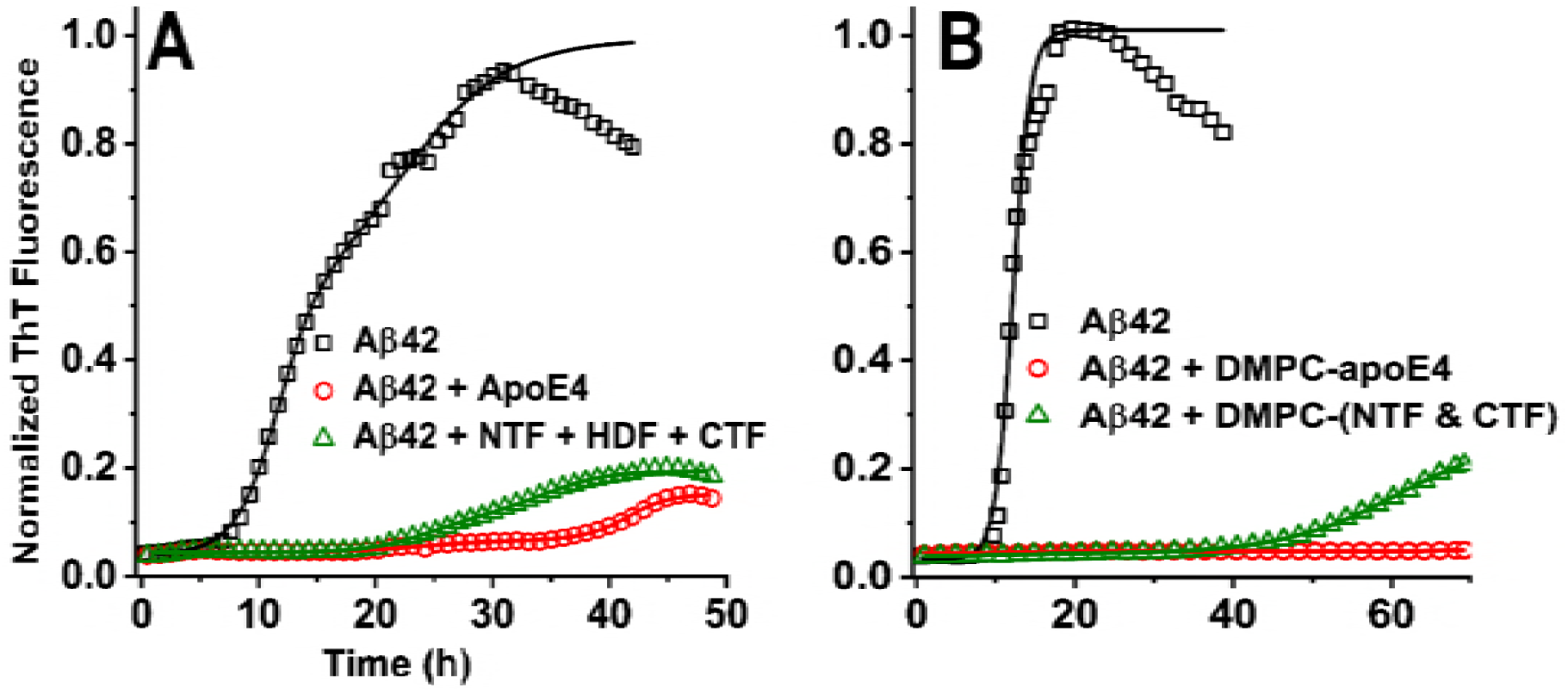
**Comparing combined effects of the fragments with full-length apoE on aggregation of Aβ.** Kinetics of aggregation of 5 μM Aβ42 in presence of 62 nM lipid-free apoE (A) and DMPC-apoE (B). Time course of aggregation Aβ42 alone (□), in presence of full-length apoE4 (○), and in presence of the fragments of apoE4 (Δ). In Panel B only DMPC-NTF and DMPC-CTF are used. The solid lines are guide to the eye. All the samples are prepared in PBS, pH 7.4 buffer and the experiments performed at 25 εC.

Therefore, NTF and CTF of apoE can independently alter the aggregation of Aβ42, and the sum of the effects of the fragments are less but comparable to the effects of the full-length apoE. Thus, both the fragments and the full-length apoE interact with Aβ with affinities in the nanomolar range.

### Effects of apoE on total aggregation monitored by fluorescence quenching of TMR-Aβ42

Since ThT fluorescence is specifically sensitive to fibril formation, the data presented above report effects of the fragments and the full-length apoE on fibril formation only. If apoE redirects aggregation of Aβ to non-fibrillar pathways it cannot be observed using fluorescence of ThT alone. Hence, we use fluorescence quenching of TMR-Aβ42 to monitor the total aggregation based on the assay developed by Garai and Frieden (35). Figure 3A-B show that kinetics of aggregation monitored by TMR fluorescence exhibits four distinct phases, an early ‘oligomerization’ phase with rapid loss of fluorescence, the ‘lag’ or intermediate phase, the growth phase and the saturation phase, consistent with the previously reported results (35). The same figures show the comparison of the combined effects of the fragments of apoE4 with that of the same molar concentration of the full-length intact apoE4. It may be seen that both the fragments and the full-length apoE4 alter the early oligomerization phase, extend the lag phase and reduces the total extent of aggregation. Furthermore, the combined effects of the fragments are comparable to the effect of the full-length apoE both in the lipid-free and lipidated forms, consistent with the observations in Figure 2. We have also verified that all the fragments of apoE can delay the aggregation of TMR-Aβ42 independently (see Figure S4), consistent with the observations in Figure 1.

**Figure 3:**
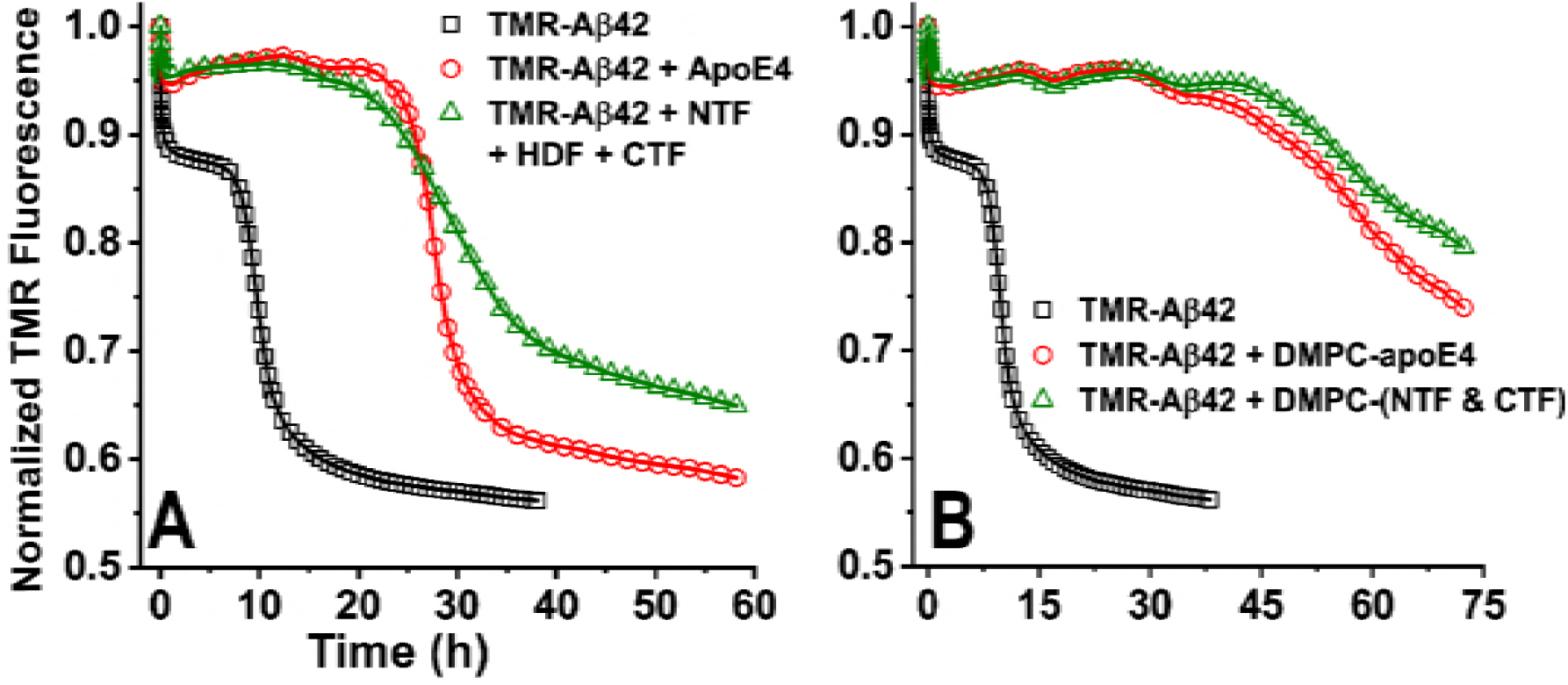
**Effects of apoE on aggregation of TMR-Aβ42**. Panel A and B show the effects of lipid-free apoE and DMPC-apoE respectively. Aggregation kinetics of 1 µM TMR-Aβ42 in absence of apoE (□), in presence of 50 nM full-length apoE (○) and fragments of apoE (Δ). In Panel B only DMPC-NTF and DMPC-CTF are used. Aggregation is monitored by quenching of TMR fluorescence with λ_ex_ = 520 nm and λ_em_ = 600 nm. The solid lines are guide to the eye. All the samples are prepared in PBS, pH 7.4 buffer and the experiments performed at 25 εC.

### Monitoring the apoE-Aβ interactions by Forster resonance energy transfer (FRET)

The data presented above clearly indicate strong interactions between apoE and Aβ. However, the literature data about the biophysical properties of the apoE-Aβ complexes are somewhat confusing. For example, in a recent review article, Tai et al have suggested that the complex formation is dependent on multiple factors such as the oligomeric status of Aβ, lipidation of apoE and the source of apoE (27). Furthermore, Verghese et al have shown that direct interactions between apoE and Aβ are minimal (5). Fluorescence correlation spectroscopy data presented here in Figure S5 indicate that apoE doesn’t interact with Aβ appreciably even at micromolar concentrations. Therefore, we hypothesize that apoE interacts weakly with the monomers of Aβ but strongly with the ‘intermediates’ or the oligomers of Aβ (11-13, 36).

To investigate the interaction between apoE and the oligomers of Aβ here we have followed the time evolution of the intermolecular FRET. In these experiments, we have used EDANS-Aβ42 and unlabeled apoE4. Fibrillization of EDANS-Aβ42 has been verified using imaging of the aggregates by AFM (see Figure S6). Here the native tryptophan residues in apoE are the FRET donor and EDANS is the acceptor. We note here that the NTF, HDF and CTF contain four, one and two tryptophan residues respectively. Figure 4A shows that all the fragments of apoE exhibit FRET with EDANS-Aβ in a time-dependent manner. The extent of the FRET is the highest for the full-length apoE4 followed by the CTF and the NTF. The FRET between Aβ and the HDF is found to be very low. The slow time-dependent increase of FRET is consistent with interactions of apoE with species of Aβ42 such as the ‘oligomers’ that form in a time-dependent manner. Therefore, the FRET data indicate that apoE may contain multiple binding sites for interacting with the oligomers of Aβ.

**Figure 4:**
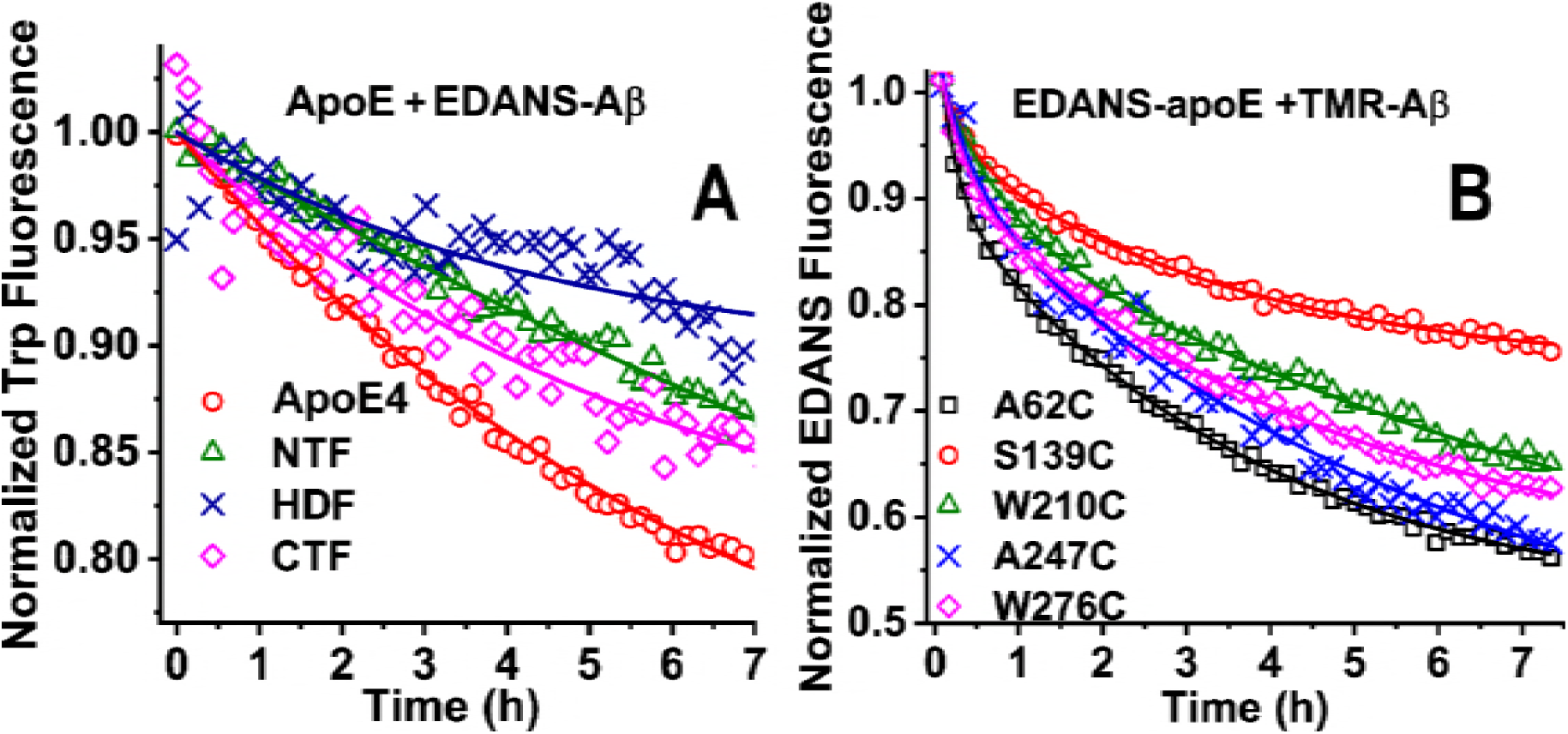
**Kinetics of interaction between ApoE and Aβ monitored by intermolecular FRET**. (A) FRET between native tryptophan residues of ApoE (500 nM) and EDANS-Aβ42 (5 µM) monitored by tryptophan fluorescence. The symbols represent experimental data using full-length apoE (○), NTF (Δ), CTF (◊) and HDF (×) of apoE. (B) Time course of FRET between 250 nM EDANS-apoE4 (donor) and 1 µM TMR-Aβ42 (acceptor) monitored by donor fluorescence. Single cysteine mutants of apoE4 are labelled with EDANS at position A62C (□), S139C (○), W210C (Δ), A247C (×), W276C (◊). The solid lines are guide to the eye. All samples are prepared in PBS, pH 7.4 buffer and experiments performed at 25 εC.

To verify the presence of multiple binding regions on apoE, further, we examined FRET between the full-length apoE labeled with EDANS (donor) and TMR-Aβ (acceptor). In these experiments, EDANS is attached to a single cysteine residue that has been introduced by site-directed mutagenesis in different locations in the sequence of apoE4. Here we have used 5 different single cysteine mutants of full-length apoE4, *viz.*, A62C, S139C, W210C, A247C, and W276C for labeling with EDANS. These mutants are chosen to cover all the three domains, *viz.*, the N- and C-terminal, and the hinge-domain. Figure 4B shows that the kinetics of FRET between apoE and Aβ are similar for all the five mutants. The extent of the FRET is the highest for the A62C region followed by A247C, W276C, W210C and S139C regions. Similar time-dependent increase of the FRET is also observed with EDANS-labeled lipidated apoE (see Figure S7). Therefore, apoE-Aβ interactions can be characterized as multivalent binding involving multiple binding sites on apoE and the oligomers of Aβ (39).

### Comparing binding affinities of the fragments and the full-length apoE using a competitive binding assay

The data presented above pose an intriguing question about whether the multivalent binding between apoE and Aβ is cooperative. The canonical way to examine this, is by comparing the binding affinities of the fragments and the full-length apoE with Aβ. In case of positive (or negative) cooperativity the binding affinity of the full-length apoE is expected to be higher (or lower) than the binding affinities of the fragments mixed together (40). However, this experiment is not straightforward here as apoE interacts predominantly with the ‘intermediates’ or the oligomers of Aβ. Since we do not know the concentration of the oligomers it is not possible to measure the binding affinities. Therefore, we attempt to compare the binding affinities qualitatively by monitoring the kinetics of FRET between EDANS-apoE and TMR-Aβ in presence or absence of unlabeled competitors, *viz.*, the unlabeled NTF, CTF and full-length apoE. Figure 5A shows that the fluorescence of EDANS labeled full-length apoE4 decreases. with time due to time-dependent increase of the FRET with TMR-Aβ, consistent with the observations reported in Figure 4B. The extent of FRET is reduced in presence of 4-fold higher molar concentration of the unlabeled NTF, CTF, mixture of NTF and CTF, and full-length apoE indicating competition between the unlabeled and the EDANS-labeled apoE proteins. However, the reduction of the FRET signal is very small in presence of the unlabeled fragments but large in presence of the full-length apoE4. Similar observations are made even in presence of mixture of all three fragments, i.e., NTF, CTF and the HDF of apoE (see Figure S8). Therefore, the fragments of apoE compete poorly with the full-length protein to interact with Aβ. Figure 5B-D show the kinetics of FRET of TMR-Aβ with EDANS-NTF + EDANS-CTF, EDANS-NTF and EDANS-CTF respectively in presence and absence of the unlabeled proteins. In all cases the FRET is reduced strongly in presence of unlabeled full-length apoE4 but affected modestly in presence of unlabeled NTF or CTF or NTF + CTF. Taken together the results presented above indicate that the binding affinities of the fragments of apoE are much weaker compared to that of the full-length protein. Interestingly, the presence of excess amount of CTF appears to enhance the FRET between the EDANS-NTF and TMR-Aβ (see Figure 5C). The reason for the enhanced FRET is not clear but this may indicate cooperativity between NTF and CTF to interact with TMR-Aβ.

**Figure 5:**
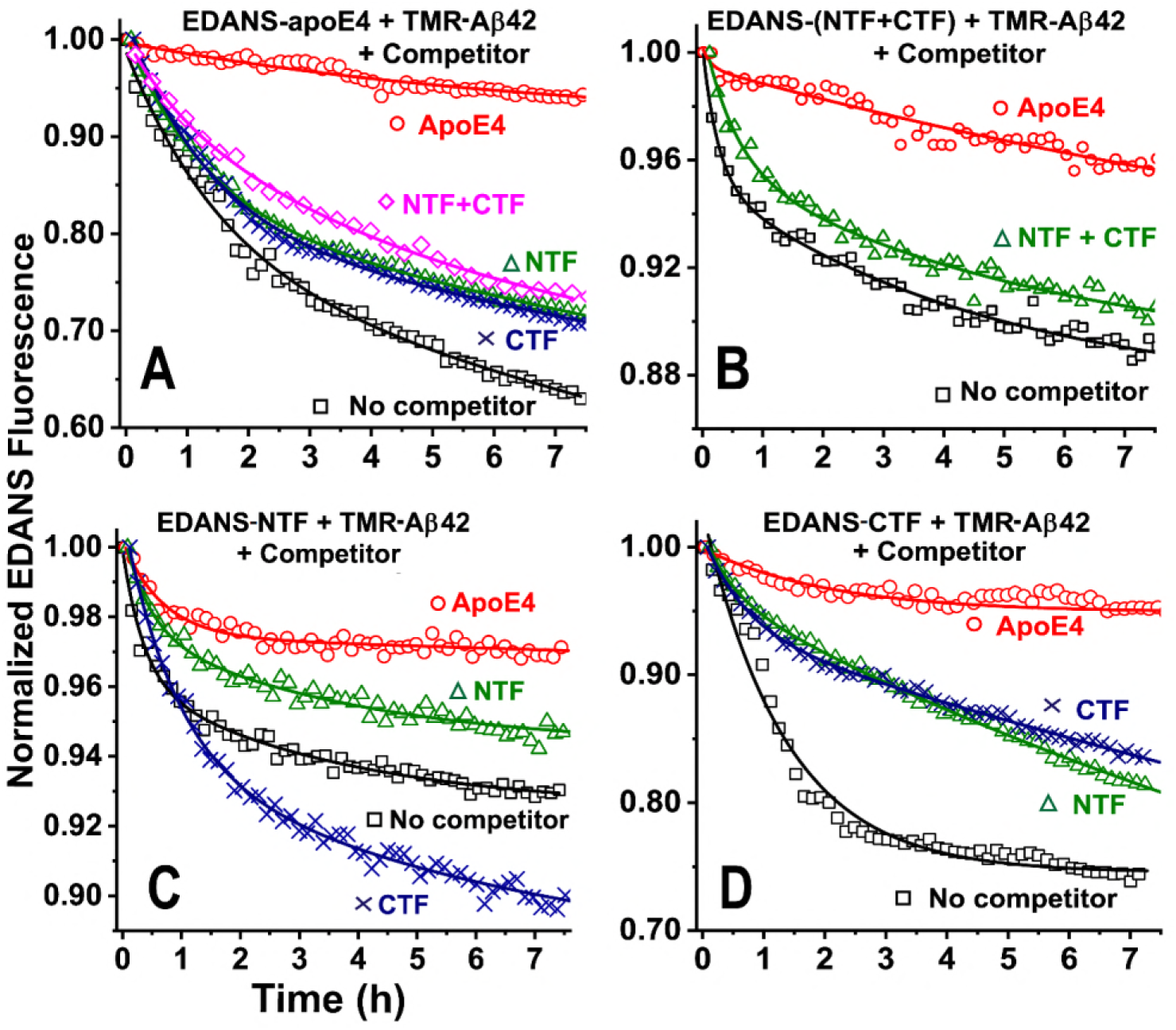
**Competitive binding monitored by FRET between EDANS-apoE and TMR-Aβ in presence of competing unlabelled apoE**. Time course of FRET between 1 µM TMR-Aβ and 250 nM EDANS-apoE4(W247C) (A), EDANS-NTF(A102C) + EDANS-CTF(A247C) (B), EDANS-NTF(A102C) (C) and EDANS-CTF(A247C) (D). The symbols represent data, in absence of competitor (□), and in presence of 1 µM unlabelled proteins, viz, full-length apoE4 (○), NTF (Δ), CTF (×) and NTF+CTF (◊). All the proteins are added at t = 0. The FRET is monitored using EDANS (donor) fluorescence. The solid lines are guide to the eye. All the samples are prepared in PBS, pH 7.4 buffer and experiments performed at 25 ^o^C.

### Reversal of FRET between EDANS-apoE and TMR-Aβ by unlabeled apoE

We examine the competition between the fragments and the full-length apoE with TMR-Aβ further by using an alternative approach. In this approach first, we monitor the kinetics of FRET between EDANS-apoE and TMR-Aβ for several hours, then we add 4-fold molar excess of the unlabeled fragments or the full-length apoE to examine if the unlabeled proteins can replace the already bound EDANS-apoE. Figure 6A shows that the fluorescence of EDANS-apoE decreases as a function of time expectedly due to FRET with the oligomers of TMR-Aβ42. Addition of excess amounts of unlabeled full-length apoE at t = 14 hr reverses the FRET in a time-dependent manner indicating reversibility of the apoE-Aβ interactions. However, the reversal of FRET in presence of the NTF and the CTF mixed together is very small consistent with weaker affinities of the fragments compared to that of the full-length protein. We note here that reversal of FRET by unlabeled full-length apoE is not 100%. This may indicate that a fraction of the apoE-Aβ complexes are highly stable. Furthermore, Figure 6B shows that the FRET between TMR-Aβ and the EDANS-NTF + EDANS-CTF can be reversed almost completely by addition of unlabeled apoE. Taken together the data presented in Figure 6 indicate that the affinity of full-length apoE4 for binding to TMR-Aβ is much higher than the resultant affinity of the fragments mixed together. It may be noted here that the reversal of FRET, i.e., the dissociation of EDANS-apoE from its complex with TMR-Aβ in presence of unlabeled apoE is a slow process taking several hours. It is possible that the apoE molecules undergo structural reorganization upon binding to Aβ oligomers forming highly stable complexes, which dissociate very slowly.

**Figure 6:**
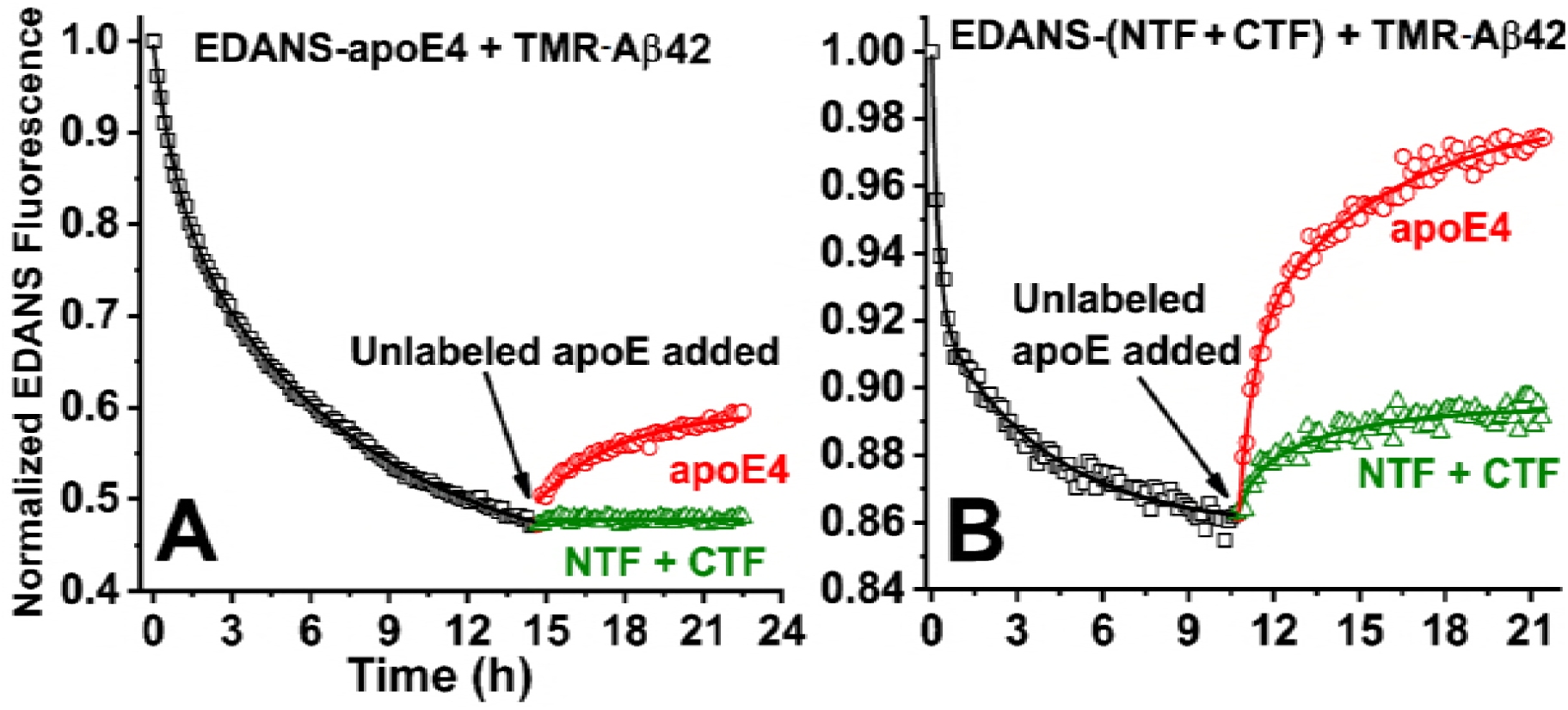
**Displacement of EDANS-apoE from EDANS-ApoE/TMR-Aβ complexes by unlabelled apoE**. FRET is monitored continuously using donor (EDANS) fluorescence from a solution of 1 µM TMR-Aβ mixed with 250 nM EDANS-apoE4(W210C) (A) or EDANS-NTF + EDANS-CTF (B) (□). Unlabelled proteins (1 µM), viz, full-length apoE (○) or NTF + CTF (Δ) is added in this sample at t > 10 hr as shown in the figures. Time dependent increase of EDANS fluorescence indicate displacement of EDANS-apoE by the unlabelled apoE from the complexes. All the samples are prepared in PBS, pH 7.4 buffer and experiments performed at 25 εC.

## Discussions

Numerous *in vivo* and *in vitro* studies indicate isoform specific role of the apoE on metabolism of the Aβ peptides in the brain (27, 41-43). ApoE-Aβ complexes have been detected both in soluble forms and insoluble plaques in the transgenic AD mice and in the human brains (7-9). Thus, understanding apoE-Aβ interaction is believed to be critically important to decipher the isoform specific roles of apoE in AD. However, biophysical properties of the apoE-Aβ complexes still remain largely unclear and somewhat confusing (reviewed in Tai et al.) (27). It is believed that the nature of the apoE-Aβ complexes are dependent on the source of both apoE and Aβ, and sometimes on the method of detection (27). The apparent confusions regarding the properties of the apoE-Aβ complexes arise likely due to the heterogeneous nature of the complexes. For example, recent biophysical studies indicate that apoE interacts with one or more intermediates of Aβ rather than the predominant monomeric forms. The ‘intermediates’ are putatively the oligomers of Aβ (11, 12, 24, 36). Therefore, the difficulties in characterization of apoE-Aβ interactions arise due to the transient and ambiguous nature of the oligomers.

### Nature of the apoE-Aβ complexes

Our results are consistent with the earlier reports, that apoE interacts with the oligomers of Aβ. First, strong inhibition of aggregation of Aβ by substoichiometric concentrations (1:160) of apoE can’t be explained by interaction with the predominant monomeric form of Aβ, rather it must interact with one or more intermediates (11-13). Second, extremely slow kinetics of the FRET between apoE and Aβ as shown in Figure 4, indicate that apoE interacts with the species of Aβ that form in a time-dependent manner. Third, FCS data presented in Figure S5 indicate no detectable interactions between apoE and Aβ even in presence of micromolar concentrations of apoE4. Therefore, high affinity interactions between apoE and Aβ doesn’t involve the predominant monomers of Aβ rather it involves ‘intermediates’ or oligomers of Aβ that form in a time-dependent manner.

The primary goal of the present work is to identify the regions of apoE that interact with Aβ. Since the interaction of apoE occurs with non-equilibrium species of Aβ, we have pursued a kinetic approach to follow the interactions between these proteins. First, we have shown that all the three fragments, *viz.*, the NTF, CTF, and HDF can delay the aggregation of Aβ even at substoichiometric concentrations, although the effects of HDF is relatively small (see Figure 1C). Then, using FRET we find that all the three domains in isolation as well as in the intact apoE interact with Aβ (Figure 4). Therefore, our data are consistent with observations by Deroo et al., who have used cross-linking followed by mass spectrometry and showed that both N- and C-terminal domains of apoE interact with Aβ (32). These observations lead us to conclude that apoE-Aβ interaction is unlike traditional enzyme-substrate interactions, which involve a specific binding site. Rather, apoE harbors multiple binding sites spread over its entire sequence.

### Multivalent interactions

Thus, apoE-Aβ interactions can be considered as multivalent binding involving multiple binding sites on the apoE molecule and the oligomeric forms of Aβ as depicted schematically in Figure 7. In the schematic, we assume that Aβ oligomers act as multivalent ligands with each Aβ molecule containing a sticky patch for self-assembly and a binding site for apoE. While the affinity of binding between individual binding sites of apoE and the monomeric form of Aβ is weak, the affinity of binding of the full-length apoE with the oligomers of Aβ could be high. Furthermore, our results show that both NTF and CTF interact with Aβ but the binding affinity of the full-length apoE is much higher. Therefore, interactions of the N- and C-terminal domains (in the intact apoE) with the Aβ oligomers is positively cooperative (see Figures 5 and 6), i.e., the free energy (ΔG_tot_) of complex formation is significantly lower than the sum of the free energies of the individual (ΔG_i_) interactions, i.e., ΔG_tot_ << ΣΔG_i_ (40). We note here that the schematic presented in Figure 7 is qualitative in nature, as the exact number of binding sites on apoE, the size of the oligomers of Aβ and ΔG of the interactions are unknown. Furthermore, in this study we haven’t examined interactions of apoE with the Aβ fibrils. However, such interactions have been reported earlier (12). We also note here that the combined effect of the fragments on the kinetics of aggregation of Aβ is found to be comparable to the full-length apoE (see Figure 2 and 3), despite the large differences in the binding affinities. The nearly equal effects on the aggregation kinetics may arise due to the following reasons. First, formation of the Aβ oligomers is slow, hence this most likely is the rate limiting process in the apoE-Aβ interactions. Second, concentrations of NTF and CTF used in our experiments are higher than the dissociation constants of the interactions of the respective fragments with the Aβ oligomers. Therefore, if the resultant binding stoichiometry of all the fragments combined is nearly equal to that of the full-length apoE then its effects on the kinetics of aggregation of Aβ could be comparable.

**Figure 7:**
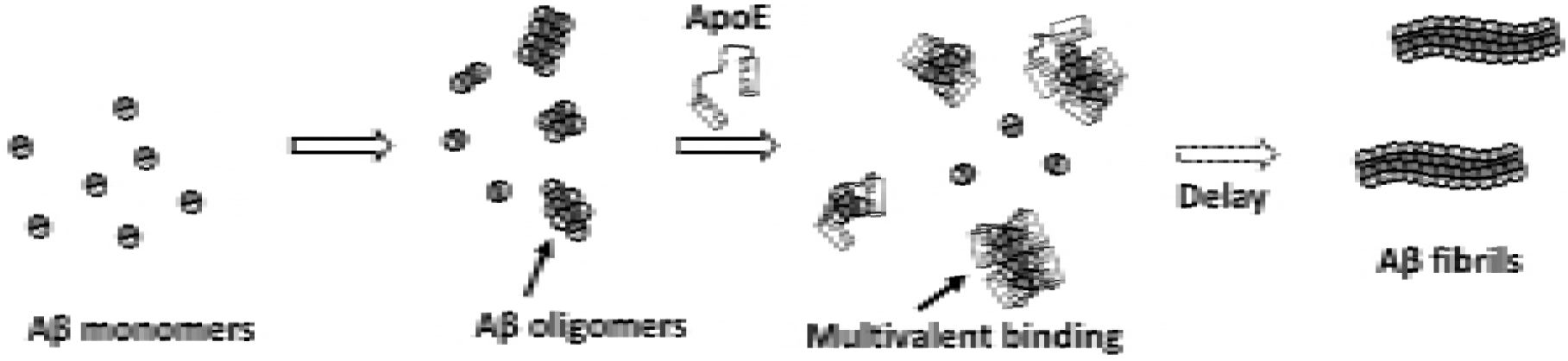
**Schematic representation of Multivalent binding of apoE with Aβ oligomers.** Aβ monomers (circles) contain at least two regions, an oligomerization motif (shaded grey) and an apoE binding motif (striped area). Thus, Aβ oligomers contain multiple patches for binding to apoE. ApoE molecules contain multiple patches for binding to Aβ (striped patches). ApoE molecules bind to Aβ oligomers via multivalent complex formation. ApoE-Aβ complexation delays fibrilization of Aβ.

Multivalent binding plays highly important roles in many biological processes such as antibody-antigen interactions, docking of cells on surfaces, cytoskeletal dynamics and transport through nucleoporin complexes etc (44-47). A growing body of evidence suggests that many of these interactions involve proteins with intrinsically disordered regions (IDRs) containing multiple short linear motifs (SLiMs) connected by flexible linkers (47). Heterotypic multivalent interactions between the SLiMs and the partner proteins are involved in the cytoskeletal dynamics and transport of macromolecules through nucleopore complexes etc (45, 47). Furthermore, this is becoming an important area of future research in developments of drugs that can bind to it targets with controllable specificity and affinity (48, 49). However, identifying and understanding the molecular details of multivalent interactions between IDRs and macromolecular binding partners have been difficult due to the ‘fuzzy’ nature of these complexes. Novel approaches including applications of single molecule techniques are required to characterize such interactions.

### Biological significance

Interactions between apoE and Aβ are believed to play critical roles in metabolism of Aβ in the brain (27, 41-43). In this work, we haven’t explored the physiological activities of the apoE-Aβ complexes. While the data presented here indicate that apoE inhibits aggregation of Aβ, physiological implications of these interactions could be several folds. For example, apoE-Aβ interactions can promote clearance, uptake, and degradation of the toxic oligomers of Aβ via the apoE receptors (27, 41). Alternatively, apoE-Aβ interactions can have harmful effects due to stabilization of the cytotoxic oligomers of Aβ (11, 12, 14, 36). Furthermore, physiological functions of apoE may be compromised due to its interactions with the Aβ oligomers (23). A major outcome of the study presented here is that high affinity interactions with the Aβ oligomers requires full-length apoE. However, proteolytic fragmentation of apoE, particularly of apoE4, have been detected in the brain around the plaques and inside the neuronal cells (29, 50, 51). Therefore, the proteolytic susceptibility of apoE4 may hinder its interaction with Aβ *in vivo* leading to inefficient clearance of the oligomers of Aβ in the brain (52). This is consistent with the reports that apoE4 is associated with impaired clearance of Aβ from ISF (4, 42).

In summary, we have clearly demonstrated that apoE interacts with Aβ and alters its aggregation *in vitro*. The apoE-Aβ interaction is high affinity and multivalent in nature. It involves multiple interactions sites located on all three domains of apoE and the oligomers of Aβ. Tight binding between apoE and Aβ is attained via positive cooperativity of the individual binding interactions.

## AUTHOR CONTRIBUTIONS

S.G. and K.G. designed research, analysed data, and wrote the paper. S.G., T.B.S. and S.D. performed research.

## Acknowledgements

The authors thank Dr. C. Neeraja for help with the expression and purification of the apoE proteins and Dr. B. Sahoo for help with Fluorescence Correlation Spectroscopy experiments. K.G. received funding from Department of Science and Technology, India.

## List of abbreviations

Aβ: Amyloid β
ApoE: Apolipoprotein E
ApoE4: Apolipoprotein E4
NTF: N-Terminal Fragment of apoE
HDF: Hinge Domain Fragment of apoE
CTF: C-Terminal Fragment of apoE
TMR: Tetramethylrhodamine
EDANS: 5-((2-Aminoethyl)amino)naphthalene-1-sulfonic acid
ThT: Thioflavin T
TCEP: (tris(2-carboxyethyl)phosphine)
βMe: β-Mercaptoethanol
EDTA: Ethylenediaminetetraacetic acid
FRET: Forster Resonance Energy Transfer

